# Superinfection exclusion creates spatially distinct influenza virus populations

**DOI:** 10.1101/2022.06.06.494939

**Authors:** Anna Sims, Laura Burgess Tornaletti, Seema Jasim, Chiara Pirillo, Ryan Devlin, Jack Hirst, Colin Loney, Joanna Wojtus, Elizabeth Sloan, Luke Thorley, Chris Boutell, Edward Roberts, Edward Hutchinson

## Abstract

Influenza viruses can interact during coinfections, allowing viral fitness to be altered by genome complementation and competition, and increasing population diversity through reassortment. However, opportunities for these interactions are limited, as coinfection is blocked shortly after primary infection by a process known as superinfection exclusion (SIE). We asked whether SIE, which occurs at the level of individual cells, could limit within-host interactions between populations of influenza viruses as they spread across regions of cells. We first created a simplified model of within-host spread by infecting monolayers of cells with two isogenic influenza A viruses, each encoding a different fluorophore, and measuring the proportion of coinfected cells. In this system SIE begins within 2-4 hours of primary infection, with the kinetics of onset defined by the dose of primary virus. We then asked how SIE controls opportunities for coinfection as viruses spread across a monolayer of cells. We observed that viruses spreading from a single coinfected focus continued to coinfect cells as they spread, as all new infections were of cells that had not yet established SIE. In contrast, viruses spreading towards each other from separately infected foci could only establish minimal regions of coinfection before SIE blocked further coinfection. This patterning was recapitulated in the lungs of infected mice and is likely to apply to other viruses that exhibit SIE. It suggests that the kinetics of SIE onset separate a spreading infection into discrete regions, within which interactions between virus populations can occur freely, and between which they are blocked.

**Importance:** Viral fitness and diversity are altered by genome interactions, which occur when multiple viruses coinfect a cell. This has been extensively studied for influenza A viruses (IAV), which use genome reassortment to adapt to new hosts and create pandemic strains, and whose replication can be compromised by the acquisition of defective-interfering RNAs. Coinfection of an individual cell by IAV is restricted by the gradual onset of superinfection exclusion (SIE). Replication of IAVs within host organisms involve the asynchronous replication of viruses as they spread to infect multiple cells. We found that under these circumstances, SIE creates spatially separated sub-populations of IAV, between which there are limited opportunities for genome interactions. Our work suggests SIE will cause many viruses to segregate into distinct subpopulations within their hosts, constraining the effects of genome interactions on their fitness and evolution.

## Introduction

Influenza viruses are a major cause of morbidity and mortality from respiratory disease, and are associated with an estimated 300,000 - 500,000 deaths globally in a typical year (1). Influenza A viruses (IAVs) are an important public health risk due both to their ability to spread as seasonal epidemics and also because they cause occasional influenza pandemics through the generation of novel influenza A virus strains (2). Influenza is able to generate new pandemic strains through reassortment, which occurs when the influenza genome segments of two or more differing influenza strains are exchanged and packaged into new virions (3). For this to occur, the two parental viruses must infect the same cell at the same time in a process known as coinfection.

Coinfection allows genome interactions to occur between viruses, which can either increase or decrease viral population fitness and diversity (4–6). An example of a genome interaction that reduces viral fitness is interference mediated by defective interfering RNAs (DI-RNAs). DI-RNAs are viral genome segments that carry a large internal deletion but which retain sequences necessary for replication and packaging into new virus particles. They replicate faster than full-length segments as they are shorter, and compete with them for incorporation into virus particles. Over multiple rounds of infection, this competition reduces the amount of infectious material within the viral population (7,8). Conversely, genome interactions can also increase viral fitness, as would occur if a virus population gains a genome segment conferring a fitness advantage (for example an antiviral escape mutation) through reassortment. Interactions between coinfecting virus genomes can also allow so called “non-infectious” influenza particles to participate in productive infection (9). A subset of these “non-infectious” particles are semi-infectious particles (SIPs), which do not contain a full set of functional viral genome segments and make up the majority of IAV populations (10). SIPs must coinfect with another particle to obtain the complete viral genome needed for a cell to produce new virus particles. Therefore, genome interactions between SIPs can result in productive infections due to ‘multiplicity reactivation’ (9–11). Therefore, coinfection allows genome interactions to occur between viruses, which is an important modulator of viral population fitness and diversity.

There are multiple lines of evidence for IAV coinfections. Viral genomics demonstrates that reassortment must occur to some degree during natural infections, implying the coinfection of cells. Most obviously, influenza pandemics have arisen repeatedly by reassortment (12), and we can also detect reassortant viruses when monitoring the viruses circulating in populations of host organisms over a period of time (13–15). In an animal model of infection, a virus that was dependent on coinfection for replication could be recovered from the guinea pig nasal passage following intranasal inoculation, indicating that coinfection can occur within this host under sufficiently strong selection (16). In carefully-controlled cell culture models of infection, reassortment can be frequent, and the proportion of reassortants increases exponentially with the frequency of co-infection (17). However, although coinfection of cells within a host organism is clearly possible, it is not been straightforward to study this directly, and the spatial context of IAV coinfections within a host is not well understood (18). Observations of discrete infectious foci have been made in the lungs of human patients and animal models infected with IAV, indicating that the viruses can spread to directly adjacent cells during infections (19–21). However, we have little information about how these foci spread and interact.

Outside of an experimental setting it is unlikely that two unrelated virus particles would reach the same cell at exactly the same moment. Instead, one would expect unrelated viruses to replicate locally within a host organism before eventually encountering the same cell. For this reason, we assume that coinfection most commonly occurs by superinfection: the infection of a previously infected cell. With many viruses the potential for superinfection is strongly limited, as following the initial infection changes occur within a cell that progressively reduce its permissivity to secondary infection. This phenomenon is known as superinfection exclusion (SIE) and has been described for numerous viruses of bacteria, animals and plants (22–27). SIE occurs within a single cell between closely genetically related viruses, and is distinct from viral interference, where the replication of one type of virus in a host suppresses the replication of another (28). For SIE, the amount of time required for a cell to become resistant to secondary infection varies depending on virus and cell type. For laboratory-adapted influenza A viruses grown in monolayers of transformed cells, the time required between primary and secondary infection for robust SIE is typically reported as around 6 hours (17,29). SIE is known to limit the potential for genome interactions between viruses in a single cell, but its impact on genome interactions as infections spread locally across multiple cells, as within a host organism, is not clear.

We wished to ask how the onset of SIE, during IAV infections of individual cells, could constrain interactions between populations of IAV spreading locally through multiple cells. To do this we needed a model system for studying genome interactions within and between locally spreading populations of IAV. Although the most striking evidence of IAV coinfections is genome reassortment, the reassortment of unrelated viral genomes as a proxy measure of coinfection is likely to underpredict the potential for coinfection between viruses, as many factors impact the ability of reassortant viruses to reassort successfully, such as the compatibility of packaging signals and the synchronicity of the viral lifecycle (30,31). We wished to use a system in which we could discount these incompatibility effects, and in which we could also model the effects of SIE on interactions between the progeny of a single infecting virus which, aside from random mutation, will be genetically identical. We therefore chose to monitor coinfection using ‘ColorFlu,’ a system of isogenic reporter viruses that differ only in the nature of a fluorophore tag fused to the NS1 protein. This approach has been previously developed for influenza viruses for use both *in vitro*, and has been used to identify coinfected cells *in vivo* following high dose intranasal inoculation of mice (32,33).

In this study we used isogenic reporter viruses to examine how the onset of SIE constrains the sites of coinfection between locally spreading populations of IAV. We observed that SIE begins early in infection and is already partially established at 2-4 hours post primary infection, therefore providing only a narrow window in which secondary infecting viruses can productively infect cells. This is a robust barrier to superinfection – we show the rate of SIE onset is mainly determined by the number of viral genomes delivered to the cell by primary infection, and that increasing the amount of secondary infecting virus has little effect on the kinetics of SIE. Using a cell culture model, we found that the kinetics of SIE onset leads to two distinct effects as viruses spread across multiple cells. Within a single coinfected focus of infection, SIE does not restrict coinfection between progeny viruses as the plaque expands. However, when two separate and growing infected regions meet, SIE restricts coinfection between their virus populations. This creates a pattern of discrete virus subpopulations, which was recapitulated in lesions in the lungs of infected mice. Therefore, our data show that SIE defines the regions where coinfection between IAVs can occur within a host, and hence controls the ability of genome interactions to shape viral fitness and evolution.

## Results

### Superinfection exclusion onset begins rapidly after primary influenza virus infection

In order to measure the degree to which SIE affects coinfection between influenza viruses, we needed to be able to distinguish non-infected, singly infected and coinfected cells. In order to do this, we used fluorescent reporter viruses (ColorFlu) (32). These viruses are derivatives of the laboratory-adapted A/Puerto Rico/8/34 (PR8; H1N1) virus, which encode a fluorophore (in this study, either mCherry or eGFP) in segment 8, which is expressed as a C-terminal fusion to the NS1 protein. As shown in figure 1A, when we used these viruses to infect Madin-Darby Canine Kidney carcinoma (MDCK) cells, the cells appeared green (eGFP only; green in figures), red (mCherry only; magenta in figures) or yellow (if coinfected; white in figures, figure 1A). The isogenic ColorFlu viruses we used should have comparable fitness. Indeed, when we monitored single cycle (Supplementary figure 1A) and multicycle (Supplementary figure 1B) growth kinetics of the ColorFlu viruses we found no significant growth advantage between eGFP and mCherry expressing viruses (Mann Whitney U Test, p>0.05). We concluded that ColorFlu viruses would be a suitable tool for modelling the onset of SIE between closely related viruses.

**Figure 1:**
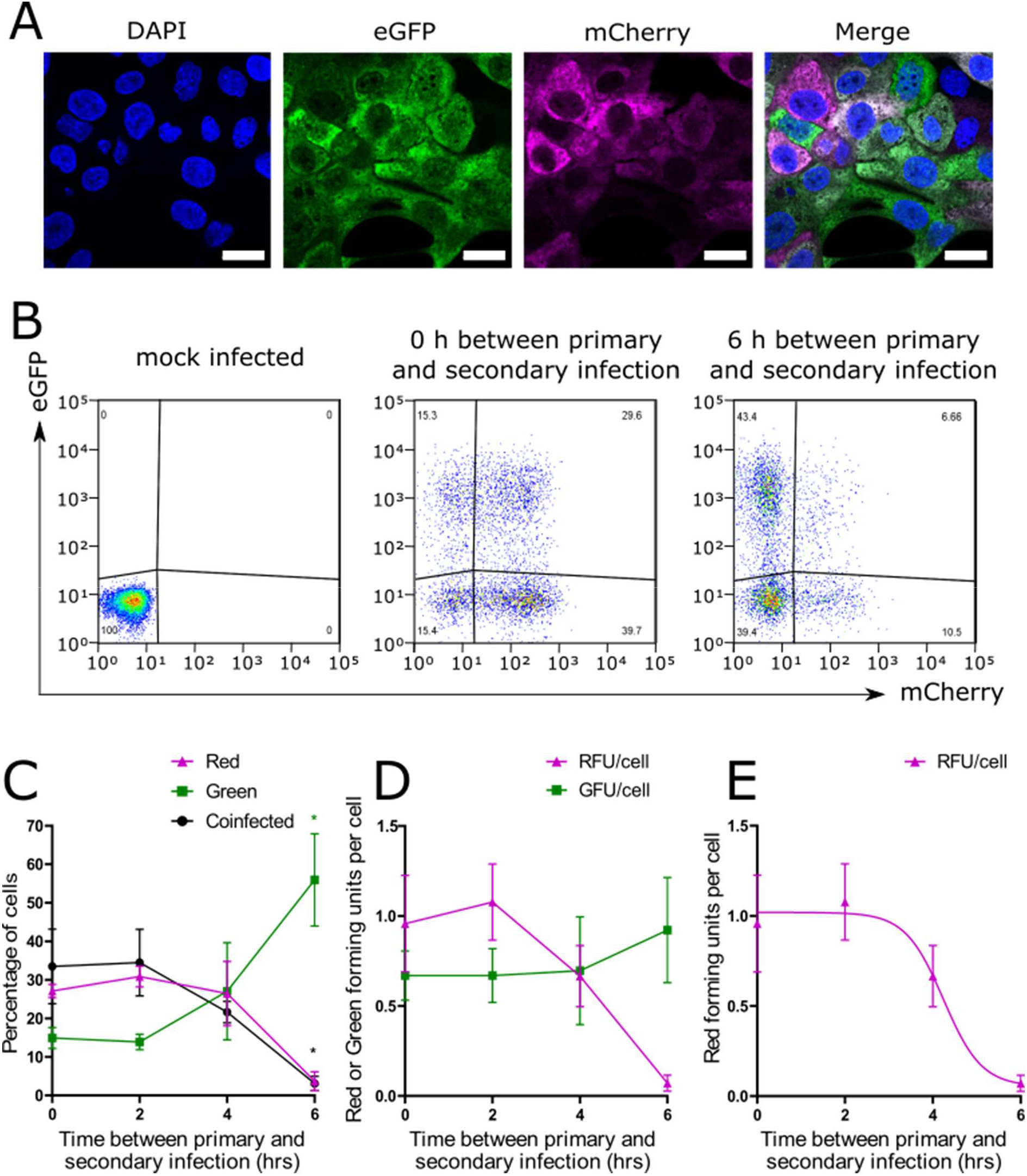
IAV induces SIE between 2-4 hours following primary infection. **(A)** Confocal micrographs of cells infected with ColorFlu viruses tagged with eGFP (green) or mCherry (magenta). Coexpression of both fluorophores is coloured white. MDCK cells were grown on glass coverslips, infected with ColorFlu viruses at MOI 0.5 for each virus, and fixed at 8 hpi. Images were obtained using a 64x objective. **(B)** Flow cytometry of cells infected with tagged viruses. MDCK cells were infected with Colorflu-eGFP before secondary infection at the time points indicated with ColorFlu-mCherry, with both viruses at MOI 1. Representative plots are shown. **(C)** Kinetics of onset of SIE, determined from flow cytometry analysis. The means and s.d. of 4 independent experiments are shown. The significance of differences from simultaneous infection were determined by Kruskal-Wallis test (* p<0.05) **(D)** The amount of red and green forming units per cell (RFU, GFU), calculated from the percentage of red, green and coinfected cells under the assumption that infection follows a Poisson distribution. The means and s.d. of 4 independent experiments are shown **(E)** A model describing an exponential reduction in RFU, fitted to data shown in part B. Total sum of squares (SST) = 0.48.

Once established, SIE blocks productive secondary infection, while prior to SIE onset cells are permissive to coinfection. In previous studies complete exclusion of secondary IAV infection has been detected by 6 hours post primary infection (34,35). We first wished to measure the onset of SIE in our model, between the time points of 0-6 hours post primary infection, and to observe the kinetics of the shift from permissivity to exclusion. To do this, we infected MDCK cells with eGFP-tagged (green) virus and then infected with mCherry-tagged (red) virus, both at an MOI of 1 FFU/cell, varying the time interval between the two infections. We then harvested the cells at 16 h after secondary infection and measured fluorophore expression by flow cytometry (figure 1B). Initially, coinfection between the viruses was unrestricted, but as the time between infection events was increased, the cells became less permissive to secondary infection, meaning that a smaller proportion could express mCherry (figure 1C). Exclusion onset in this system occurred when superinfection was between 2-4 h after primary infection, and a significant reduction in the proportion of coinfected cells (p=0.045, Kruskal-Wallis test) was detectable by 6 hpi (figure 1C). Our data are consistent with previous studies in which exclusion of the first virus was detectable if superinfection occurred 6 hours post primary infection, and we additionally show a progressive shift from a permissive to exclusionary state beginning around 2h post primary infection.

To further investigate the kinetics of SIE onset, we wanted to consider how SIE affected the ability of the primary (green) and secondary (red) viruses to infect cells. By measuring the proportion of fluorescent cells and inferring what proportion had lost the expression of the red fluorophore, we were able to quantify the extent of SIE. To do this we calculated the concentrations of viruses capable of causing cells to become red or green (‘red forming units’ (RFU) per cell and ‘green forming units’ (GFU) per cell, respectively), by assuming that simultaneously-administered viruses were able to infect cells independently of each other and that infection could therefore be modelled by a Poisson distribution. We found the GFU per cell remained stable as the interval between primary (green) and secondary (red) infection was increased. However, the RFU per cell decreased rapidly after 2 hours, showing that the red virus was excluded from the cells (figure 1D).

The mechanism for SIE in IAVs is not yet known, though it has previously been suggested that it may require an actively replicating influenza polymerase (35). We reasoned that, as the products of viral transcription and replication appear to accumulate exponentially in a newly infected cell (36,37), the inhibitory factor that drives SIE might also increase exponentially following primary infection. If this were the case, we would expect the changes in SIE to be a close fit to a model describing an exponential reduction in the RFU per cell. To test this hypothesis, we fitted a log(inhibitor) model to the RFU per cell over time (figure 1E). The model was a good fit to the data (total sum of squares (SST) =0.48), making an exponentially increasing inhibitory factor a plausible explanation for the kinetics of SIE in IAV (38).

### The main viral determinant of superinfection exclusion kinetics is the amount of primary infecting influenza virus

Once we could model the kinetics of SIE onset, we were able to examine the kinetics of SIE by asking how changing the conditions of infection affected the model parameters. Under our model, the rate of SIE onset could be assessed by the time between infections required for the maximum RFU per cell to reduce by half (the time to IC50), which in our initial experiments was 4.24 (± 0.43) hours.

We first asked if we could speed up the onset of SIE by increasing the dose of primary virus. Therefore, we repeated our measurements of SIE kinetics, increasing the input of the primary (green) virus from a baseline MOI of 1 by either 2.5 or 5 fold (figure 2A) and determining the time to IC50 of SIE (figures 3 B, C). The exponential model was a good fit to the data under all conditions tested (SST ≤ 1). When the amount of primary infecting virus (ColorFlu-eGFP) was increased, the time to IC50 of SIE decreased (figure 2B). For a 5-fold increase of the primary virus over the secondary infecting virus (ColorFlu-mCherry) this difference was statistically significant (p<0.05, Kruskal-Wallis test; figure 3B), indicating that SIE kinetics are sensitive to the amount of primary input virus.

**Figure 2:**
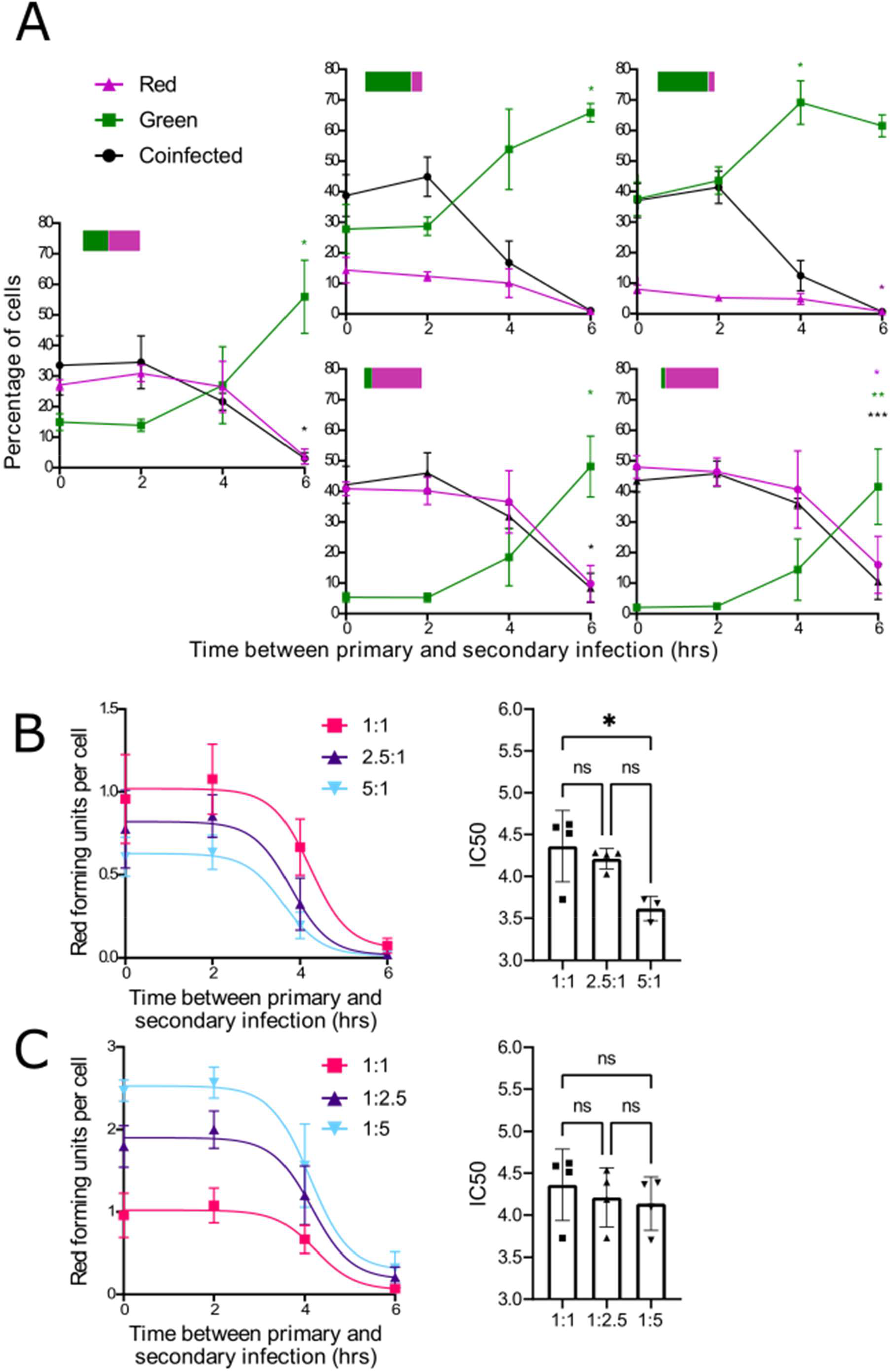
SIE kinetics are sensitive to the amount of primary infecting genomes but not the amount of secondary infecting genomes. **(A)** The effect of altering the ratios of primary (ColorFlu-eGFP, green) or secondary (ColorFlu-mCherry, magenta) viruses on coinfection. MDCK cells were infected and analysed by flow cytometry as in Figure 1, the ratios of viruses in each case are indicated as bars. The means and s.d. of 4 independent experiments are shown. The significance of differences from simultaneous infection was assessed using the Kruskal-Wallis test (*p<0.05, **P<0.01, ***p< 0.001) **(B, C)** The effect of altering the ratios of primary **(B)** and secondary **(C)** viruses on SIE. The expression of secondary virus (RFU) at the ratios shown was inferred from the data in A and fitted to an exponential reduction model as in Figure 1. (Total sum of squares (SST) for 1:1, 2.5:1, 5:1 are 0.48, 0.31 and 0.064 respectively; and for 1:1, 1:2.5 and 1:5 are 0.48, 0.88 and 1.00 respectively.) For each experiment the time between primary and secondary infection required for 50% inhibition was calculated (IC50). RFU and inhibition times are shown, with the means and s.d. of 4 independent experiments; for inhibition times the significance of differences from a 1:1 ratio were determined by Kruskal-Wallis test (*p<0.05).

**Figure 3:**
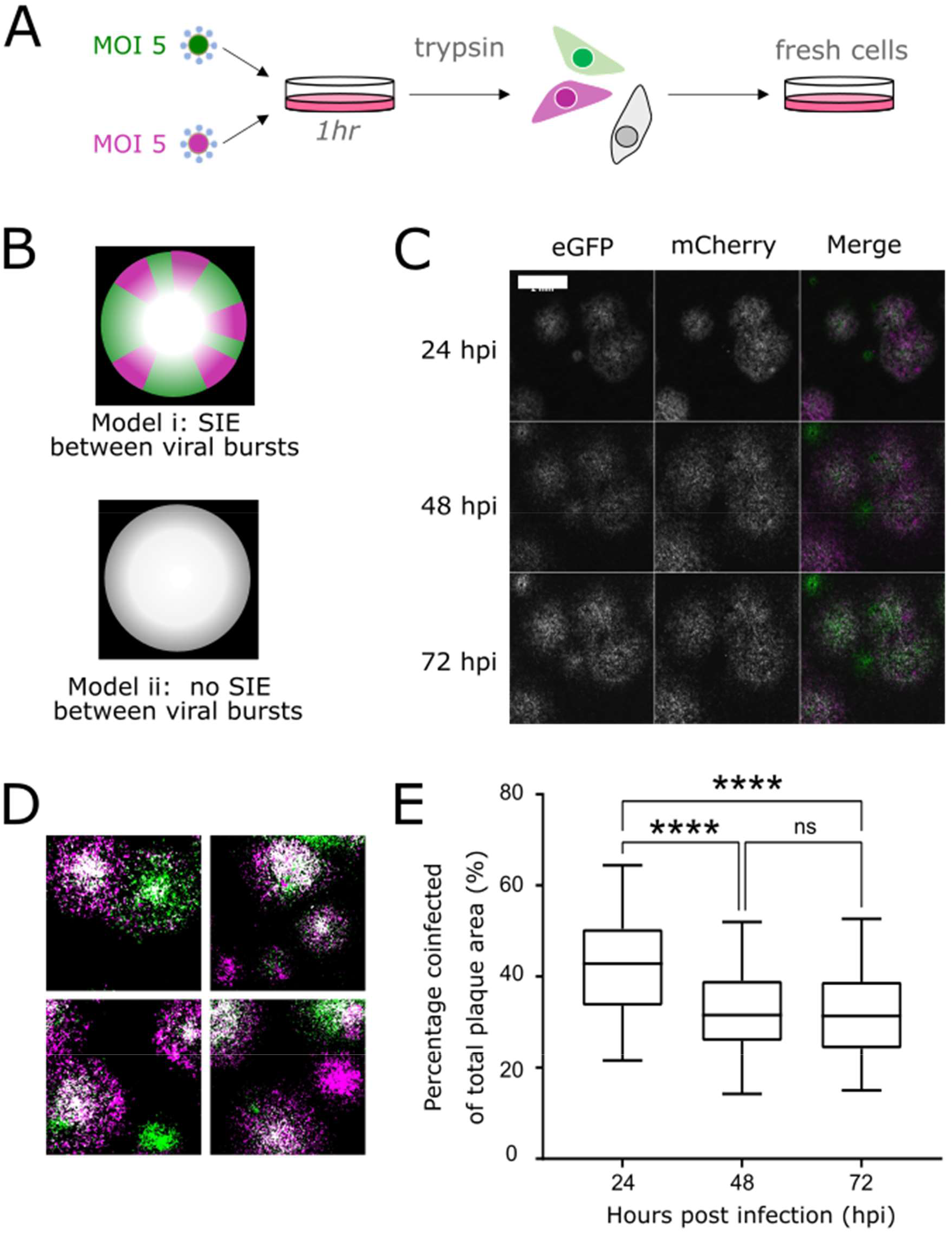
SIE does not inhibit coinfection between IAVs from a single focus of infection. **(A)** Experimental design for investigating the role of SIE in the spread of coinfected foci. **(B)** Proposed models for the spread of coinfected foci. **(C)** Representative image of coinfected plaque spread. Viruses were seeded onto monolayers of MDCK cells, overlayed with agarose and imaged every 24 h. Images were taken on Celigo fluorescent microscope. Scale bar = 2mm **(D)** Representative images at 48 hpi with applied binary threshold to distinguish coinfected cells (white) from singly infected cells (magenta or green) **(E)** Percentage of coinfected areas in comparison to total infected area, calculated from images at taken at each time point. Box and whisker plots show the percentage areas from individual fields of view (n=71) from one experiment. The significance of differences between timepoints was tested by One-Way ANOVA (**** P<0.0001).

We reasoned that the primary infecting virus might exclude a superinfecting virus through direct competition, for example by binding to cellular factors or occupying a subcellular niche (36,37). This would be consistent with observations that within 2-4 hours of primary infection the amount of viral RNA within an infected cell rises dramatically (39). If SIE is a form of competitive inhibition, we reasoned that we might be able to partially overcome SIE by increasing the amount of secondary infecting viral genomes entering the cell. When we increased the secondary (red) virus by 2.5 or 5-fold, our exponential model remained a good fit to the data (SST ≤ 1). However, we did not detect any significant shift in IC50 value at the ratios we examined (p<0.05, Kruskal-Wallis test, figure 2C). This shows that, within the range of MOIs tested, SIE kinetics are fairly insensitive to the amount of secondary infecting virus.

When taken together, our data suggest that the amount of primary infecting virus sets the kinetics for the onset of SIE and the secondary infecting virus has limited ability to overcome this. This suggests that, once established, the switch from a permissive to exclusionary state in the cells is extremely hard to overcome.

### Superinfection exclusion does not restrict interactions between influenza viruses within a spreading infection

After we defined the time frame for SIE onset, we wanted to investigate whether SIE prevents the progeny viruses produced from an initial infection from interacting with each other as an infected focus expands. To examine this we set up plaque assays, allowing viruses to propagate through MDCK cells under agarose, as a simplified model of the foci of infection observed in infected patients (21). To study interactions between the progeny of a single infected cell, we first infected MDCK cells with a mixture of green and red viruses, both at an MOI of 5, to create a population of coinfected cells. At 1 h post-infection, before new virus particles were produced, we dispersed these infected cells using trypsin, diluted them, and then applied them to a fresh MDCK monolayer and overlaid with agarose, so that each coinfected cell would be an individual plaque forming unit shedding both red and green virus (experimental procedure in figure 3A).

We hypothesised two possibilities for how the viral progeny of these cells would interact to produce plaques, either (i) rapid SIE onset would inhibit coinfection, resulting in the initial yellow focus segregating into discrete regions where one fluorophore would dominate, or (ii) SIE would not develop quickly enough to prevent coinfections at the plaque edge, and so the plaque would remain coinfected as it expands (figure 3B). We observed that as coinfected plaques expanded, both fluorophores were expressed across the entire plaque area (figure 3C). Therefore, we concluded that the cells at the leading edge of the plaque were receiving multiple viruses quickly enough for coinfection to occur before the effective onset of SIE.

Although both mCherry and eGFP expression could be detected across the plaques, we did notice that coinfected cells were concentrated towards the middle of the plaque area (figure 3D). To quantify this, we measured the areas of individual plaques that were coinfected compared to their total area. We found the coinfected portion of plaques is at its highest at 24hpi and is significantly reduced in the larger plaques that form by both 48hpi and 72hpi (figure 3E, p<0.0001 in both cases). This change in the distribution of fluorescent cells may not be due to changes in SIE however, as live-cell imaging showed infected cells migrating to the centre of plaques, presumably as they began to die (Supplementary Movie 1). Taking our data together, we concluded that the kinetics of SIE allow coinfection to occur freely between the progeny viruses from a focus of infection.

### Superinfection exclusion strongly inhibits coinfection between established regions of influenza virus infection

Next, we wanted to assess whether coinfection was restricted when viruses from two separate foci of infection expand and interact with each other. Previous studies have shown that IAV reassortment can be readily detected both *in vivo* and *in vitro* (17,40). This indicates that coinfection between different strains is possible despite the restrictions of SIE but it does not give us information as to how this occurs in the spatial context of influenza viruses as they spread. In a natural infection we assume that it is unlikely that multiple ‘incoming’ viruses would reach the same cell within a short space of time. Instead, we assume that coinfection of cells with different strains of virus typically occurs through interactions between the progeny of separate foci of infection. To model the interactions between spreading foci of infection, we infected MDCK monolayers at a low MOI with green and red viruses, overlaid with agarose and imaged the spread of plaques every 24 h for 72 h. We observed that, as adjacent plaques expressing different fluorophores grew towards each other, regions of cells expressing different fluorophores remained almost entirely distinct (figure 4A, further examples in supplementary data 2). On close examination, we observed a very thin boundary region of cells in which both fluorophores were expressed (figure 4B). Image analysis showed that the coinfected region at 72hpi was around 1% of the total plaque area (figure 4C). This indicates that only a small region of coinfection was possible before further interactions were blocked by the onset of SIE.

**Figure 4:**
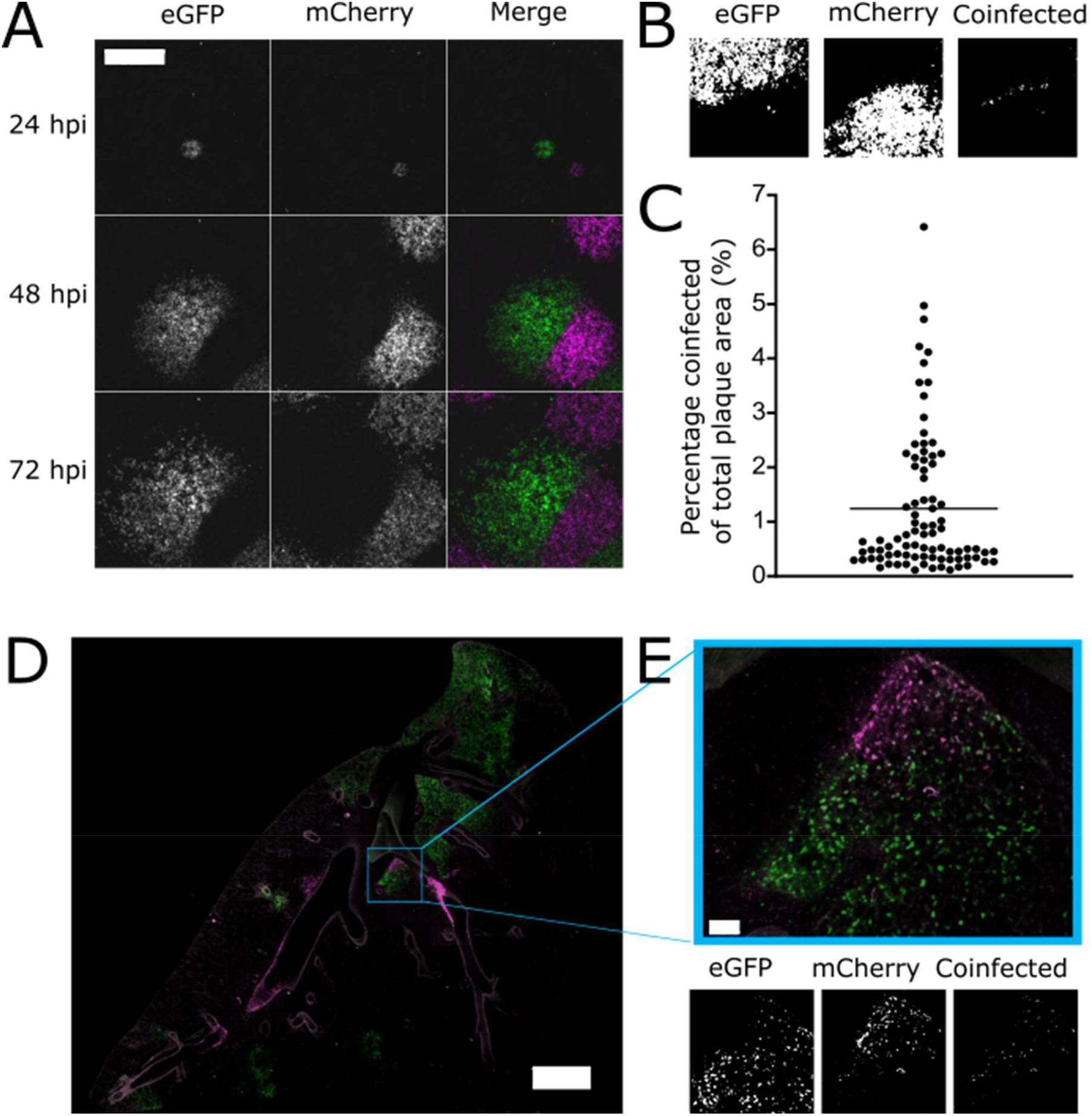
SIE allows only a small region of coinfection when two established regions of IAV infection meet. **(A)** Representative image of plaque interaction. Viruses were seeded onto monolayers of MDCK cells, overlayed with agarose and imaged every 24 hours. Images taken on Celigo fluorescent microscope. Scale bar = 2mm **(B)** Representative plaque with coinfected region highlighted. Images were applied with a binary threshold and the pixels where both red and green fluorescence were generated into a separate image displaying coinfection **(C)** Percentage of coinfected areas in comparison to total plaque area was calculated from images at taken at 72hpi. Each point represents the percentage area from a single field of view (n=86) from one experiment and the line represent the mean. **(D)** Images of lung sections from infected mice 6dpi. B57BL/6 mice were intranasally inoculated with mixtures of mCherry and eGFP expressing viruses (500 pfu of each virus). Lung sections at 6 days post infection were imaged using [confocal microscope] using 20x objective lens. Whole lung image, scale bar = 1500µm, for enlarged image of indicated area, scale bar = 100µm. **(E)** Image of representative lesion with coinfected region highlighted. Images were processed as described in (B).

To investigate whether this exclusion phenotype was relevant to infections *in vivo*, we performed a version of this experiment in which IAV spreads through the lung of a mouse. To do this, we infected C57BL/6 mice intranasally with a mixture of Colorflu-eGFP and ColorFlu-mCherry (500 PFU of each virus), took sections of lungs from mice at day 3 or 6 post-infection, and looked for regions where red and green foci of infection were interacting. Images of lungs harvested at day 3 suggested that the initial sites of infection were mainly in the bronchi (supplementary data 3). Consistent with previous reports (32), coinfection was visible at many of these sites, which presumably received a high dose of both viruses simultaneously. However, by day 6 infection had spread into the alveoli and established distinct red and green lesions. At this time point we observed multiple instances where red and green lesions were adjacent to each other but maintained a distinct boundary, despite a lack of obvious anatomical compartmentalisation (figure 4D, additional examples in supplementary data 4). This recapitulates the phenotype we observed in cell culture, and indicates that the spatial segregation of viral subpopulations we observed in that reductionist model also occurs during the propagation of viruses within a host organism. We therefore concluded that, when two initially separate regions of influenza virus infection spread and contact each other, the kinetics of SIE ensure that the potential for coinfection between these viral populations is severely inhibited.

## Discussion

SIE has been observed for many different viruses, whose hosts include plants, bacteria and animals (22–27). Although SIE is a widespread property of viruses it can be achieved through many different mechanisms, and its implications for the evolution of medically important viruses, such as the influenza viruses, is not well understood. SIE constrains the ability of related viruses to coinfect cells, as can occur when viruses replicate and spread locally through cells within a host organism. The context of this stage in viral replication – between an initiating infection of an individual cell and the transmission of replicated virus to new host organisms – is usually sequestered inside the host organism and is challenging to study directly (18). Here, we used a simplified cell culture model of infection using isogenic, fluorescently tagged IAV to derive a model for how the kinetics of SIE onset limit coinfection during spreading IAV infections, and then showed that the patterns of infection this predicts are recapitulated in experimental infection of the mouse respiratory tract. We show that that during the local spread of IAV infection from cell to cell, SIE defines the regions where coinfection can and cannot occur. Our data show that the kinetics of SIE onset in a spreading infection allow ongoing genetic interactions between the viral progeny of a single infected focus, but strongly inhibit genetic interactions between viruses from distinct foci of infection.

IAVs have come to be seen as particularly capable of coinfection, in part because of the importance of reassortment in generating new pandemic strains of IAV through ‘antigenic shift.’ In addition to inferences from the natural evolution of IAV, reassortment of IAV during coinfection can be demonstrated experimentally both *in vivo* and *in vitro* (17,40). These observations contrast strikingly with our own data, which show that SIE should impose severe restrictions on coinfection between viruses that propagate locally within a host. There could are several plausible explanations for why reassortant IAVs are regularly detected despite the effects of within-host SIE. Firstly, when considering epidemiological evidence for reassortment, the number of host organisms infected by IAVs is extremely large (38,41), providing ample opportunities even for rare interactions between viral strains within a host. Secondly, when considering experimental studies of reassortment and coinfection within animals, the studies are often designed using high concentrations of viruses administered by artificial routes such as intranasal inoculation of mammals or injection into embryonated chicken eggs (32,42,43). The delivery of a high concentration of viruses in a small window of time to the same anatomical site would be expected to increase the likelihood of coinfection of cells during an initial infection, when compared to natural infections. Our data are compatible with coinfection occurring naturally as a rare event, but indicate that the interaction of SIE with the spatial dynamics of virus spread will establish previously unstudied barriers to reassortment in natural infections.

Although we found that interactions between viruses from separate established infections were strongly inhibited by SIE, we also found that the progeny of a single parental virus infection were free to interact with each other through coinfections, unhindered by the onset of SIE. While this does not allow the reassortment of genomes from different viral strains, it could still have implications for the fitness of the population. Namely, it allows the semi-infectious progeny virions which make up the majority of the viral population (10) to complement each other. Coinfection allows these otherwise “non-infectious” virus particles to contribute to productive infections (44,45).Stochastic simulations in bacteriophage have demonstrated that viral populations that are incapable of initiating SIE are more able to fix beneficial mutations (46). Therefore, unrestricted coinfection between the progeny of a virus could help to maintain viral population fitness. However, it would also leave the viral population vulnerable to interference mediated by DI-RNAs. This is especially true as the surrounding cells receive many hundreds of virions as the plaque expands, and high MOI infections increase the likelihood of DI-RNA generation (5,47). Although this could lead to an individual lesion being overtaken by DI-RNAs, our model suggests that DI-RNAs generated in one lesion would not be able to overtake the viruses in a separate adjoining lesion due to established SIE preventing coinfection.

The results from our model confirm previous findings that SIE restricts coinfections of influenza viruses that occur after 6 h of primary infection (29,35,48). Furthermore, they show that SIE becomes detectable between 2 and 4 h after primary infection and then becomes increasingly effective. A number of models could be consistent with these data, but a model that proposes an exponentially increasing inhibitory factor is consistent with the exponential accumulation of viral products from the primary infecting virus (39), and is therefore consistent with previous observations indicating a connection between SIE and the presence of replicating influenza polymerase complexes (35). This exponential inhibition model is also consistent with previous work showing that more coinfected cells could be detected if fewer replication-competent genome segments were delivered during primary infection (35), and with our data showing that the amount of primary infecting genomes determines the kinetics of SIE onset, as exponential relationships are sensitive to their starting parameters. More work is required to determine if the mechanism of IAV SIE is due directly to the accumulation of products of replicating polymerases (either RNA transcripts or, indirectly, viral proteins), to a host-encoded factor that is produced in response to polymerase activity, or to a combination of effects.

Importantly, our model of the effects of SIE in a locally spreading IAV infection relies on the kinetics of SIE onset, rather than on a specific mechanism. It should therefore be generalisable to the large number of other viruses that establish SIE gradually during the infection of a cell, propagate locally within a host, and whose fitness and evolution are shaped by genetic exchange between viruses during coinfection. Our data imply that within a host’s tissues, at a scale between the well-studied extremes of an individual cell and a population of host organisms, viruses will naturally segregate out into a complex microscopic landscape of subpopulations, whose genetic interactions are controlled by SIE.

## Materials and Methods

### Cells and Viruses

Madin-Darby Canine Kidney (MDCK) cells (a gift from Prof. P Digard at the Roslin Institute, University of Edinburgh) and human embryonic kidney (HEK) 293T cells (a gift from Prof. S Wilson, MRC-University of Glasgow Centre for Virus Research) were maintained in complete media (Dulbecco’s Modified Eagle Medium (DMEM, Gibco) supplemented with 10% Foetal Bovine Serum (FBS, Gibco)). All cells were maintained at 37°C and 5% CO_2_ in a humidified incubator.

The wild-type (WT) PR8 was generated in HEK293T cells using the pDUAL reverse genetics system, a gift of Prof Ron Fouchier (Erasmus MC Rotterdam); as previously described (34). ColorFlu viruses (ColorFlu-eGFP and ColorFlu-mCherry) were rescued in HEK293T cells from plasmids encoding the NS segment supplied by Prof. Y. Kaowaoka (University of Wisconsin-Madison, University of Tokyo), in addition to WT PR8 pDUAL plasmids edited to contain the compensatory mutations (HA T380A and PB2 E712D) as previously described (22). The viruses were then passaged at low MOI in viral growth media (VGM) (DMEM with 0.14% (w/v) bovine serum albumin (BSA) and 1µg/µl TPCK-treated trypsin) to create a working stock.

Virus plaque titres in plaque forming units per mL (PFU/mL) were obtained in MDCK cells under agarose, following the procedure of Gaush and Smith (49).

### Mouse Infections

C57BL/6 mice (Charles River, UK) were infected intranasally with a total of 1000 PFU of ColorFlu viruses (an equal mixture of mCherry and eGFP variants). All animal work was carried out in line with the EU Directive 2010/63/eu and Animal (Scientific Procedures) Act 1986, under a project licence P72BA642F, and was approved by the University of Glasgow Animal Welfare and Ethics Review Board. Animals were housed in a barriered facility proactive in environmental enrichment.

### Immunofluorescence and Imaging

Confocal images of infected cells were obtained by infecting cells on coverslips, with an MOI of 0.5 (based on plaque titre) for each of the ColorFlu viruses, for 8 hours before fixation in 4% (v/v) formaldehyde diluted in PBS (Sigma). Following fixation, the cells were rinsed in PBS and the nucleus stained with 4′,6-diamidino-2-phenylindole (DAPI, ThermoFisher). Coverslips were then mounted and imaged with the Zeiss Laser Scanning 710 confocal microscope images were processed using Zeiss Zen 2011 software.

To obtain images of viruses spreading from a coinfected focus, MDCK cell monolayers were infected with mCherry and eGFP tagged viruses both at an MOI of 5. At 1h p.i., the infected cells were dispersed with TrypLE express for 15 minutes (ThermoFisher) and diluted in VGM to create a suspension that was applied to fresh MDCK cell monolayers. The cells were left to settle for 4 h, after which an agarose overlay was added and infections were left to proceed, as in a standard plaque assay.

To obtain images of interactions between initially separate foci of infection, MDCK cell monolayers were infected with a diluted mixture of mCherry and eGFP tagged ColorFlu viruses after which an agarose overlay was applied and infections were left to proceed as in a standard plaque assay. The infected plates were imaged through the agarose every 24 hours in a Celigo imaging cytometer (Nexcelom). Images were processed in FIJII ImageJ (50) using custom macros which can be accessed here: https://github.com/annasimsbiol/colorflu

To obtain live cell images of spreading infections, ColorFlu-eGFP and mCherry viruses were diluted in VGM to an MOI of 0.5 (PFU/mL) and applied to confluent MDCK cell monolayers. Following a 1 h incubation the inoculum was removed and agarose was overlaid, as in a standard plaque assay procedure (1). The plate was transferred to an Observer Z1 live-cell imaging microscope (Zeiss, USA), and a tile from a well was imaged every 15 mins over 72h. The acquired videos were compiled using Zen (Zeiss).

To obtain images of infections in mice, at the indicated number of days post infection animals were sacrificed and their lungs inflated with 2% low melt agarose. Lungs were then fixed in PLP buffer (0.075 M lysine, 0.37 M sodium phosphate (pH 7.2), 2% formaldehyde, and 0.01 M NaIO_4_) overnight, 300 µm sections of lung were cut using a vibrotome, and imaging was performed using an LSM 880 confocal microscope (Zeiss) using a 20x objective at 0.6x digital zoom with 5 µm z steps. Images were stitched and a maximum intensity projections were made using Imaris software (version 9.7.0, Bitplane, USA).

### Viral Growth Kinetics

For single cycle growth kinetics, viruses were applied to confluent MDCK monolayers at an MOI of 2.5 and the cells were incubated with the inoculum for 1 h at 37°C and 5% CO_2_ in a humidified incubator to allow the viruses to enter cells. Following this, the inoculum was removed, and the cells bathed in acid wash (10mM HCL and 150mM NaCl in MiliQ-water, pH3) for 1 minute after which fresh VGM was added. Media were sampled at the time points indicated, clarified by low-speed centrifugation and stored at −80°C before titration by plaque assay.

Multicycle kinetics were determined as above, except that the cells were infected at an MOI of 0.001 and the acid wash step was omitted.

### Flow cytometry

MDCK cells were inoculated for 1 hour with ColorFlu-eGFP viruses diluted in VGM at the MOI indicated. After 1 hour the inoculum was removed and replaced with complete media. After the time intervals indicated, cells were inoculated for 1 h with Colorflu-mCherry, at the MOI indicated. After 1 h the inoculum was removed and replaced with complete media, and the cells were incubated for a further 16 h at 37 °C. The proportions of cells expressing the different fluorophores were assessed using a Guava easyCyte HT System cytometer (Luminex). Briefly, infected and mock-infected MDCK monolayers were dissociated TrypLE express for 15 minutes (ThermoFisher) and dispersed into a single-cell suspension before fixation in 2% formaldehyde (v/v) in PBS. Each sample was prepared in technical triplicate and the data were analysed in FlowJo software v10.6. The thresholds for assessing positive detection of either red or green fluorophore was set using the mock-infected cells as a negative control.

### Modelling

The MOI of viruses that could cause fluorescence in different channels (red forming units (RFU) and green forming units (GFU) per cell) was calculated under the assumptions that viruses that are added to cells at the same time infect independently of each other and that all cells are equally susceptible to infection, meaning that the number of viruses infecting each cell follows a Poisson distribution (51). Under these assumptions, the mean number of fluorescent forming units (FFU) (of a particular colour) infecting a cell is given by *– ln (1 – (F+ C))*, where *F* is the proportion of cells that only express the fluorophore of interest and *C* is the proportion of coinfected that express both fluorophores. The change in RFU per cell with increasing intervals between primary and secondary infection was fitted to a model describing an exponential decrease using a four-parameter logistic curve (log(inhibitior) vs response model with variable slope) using GraphPad Prism (version 9; GraphPad).

## Acknowledgements

We acknowledge funding from the UK Medical Research Council (MRC) as studentships to A.S. [MC_ST_CVR_2019] and J.W. [MC_ST_U18018], as CVR core funding to C.B. [MC_UU_12014/5] and as a Career Development Award and Transition Support Award to E.H. [MR/N008618/1 and MR/V035789/1]; funding from the University of Glasgow to E.H., and funding from Cancer Research UK (CRUK) to E.R. [A_BICR_1920_Roberts]. The authors declare no conflicts of interest.

## Author Contributions

A.S.: conceptualisation, visualisation, methodology, investigation, writing - original draft and presentation, L.BT.: conceptualisation, visualisation, methodology, investigation S.J.: conceptualisation, visualisation, methodology, investigation C.P.: methodology, investigation R.D.: methodology, investigation J.H.: conceptualisation, visualisation, methodology, C.L.: visualisation, methodology, J.W.: conceptualisation, methodology, E.S.: conceptualisation, methodology, L.T.: conceptualisation, methodology, C.B.: supervision, E.R.: methodology, investigation, visualisation, E.H.: conceptualisation, methodology, supervision, writing - review and editing.

## Supporting Information

**SUPPLEMENTARY DATA 1:**
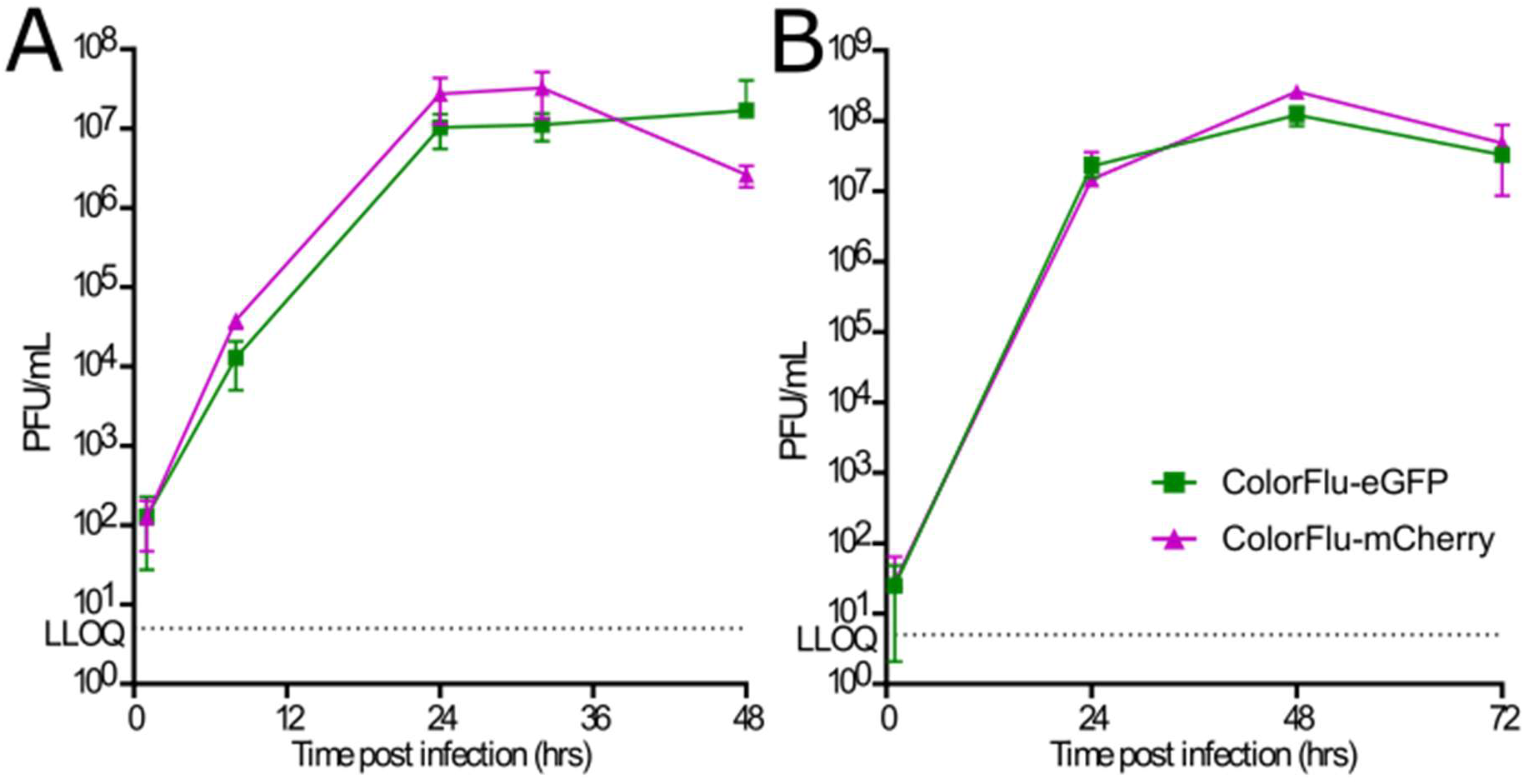
ColorFlu viruses is a suitable tool for modelling coinfection between related viruses. **(A)** Single cycle growth kinetics of ColorFlu viruses was assessed by infecting MDCK cell monolayers at MOI 2.5 and the supernatant harvested at the time points indicated. Virus titre was assessed using plaque assay on MDCK cells. **(B)** Multi-cycle growth kinetics of ColorFlu viruses was assessed by infecting MDCK cell monolayers at an MOI of 0.001 and sampled as described in single-cycle growth kinetic assay. For both B and C, values represent the mean + SD for three independent experiments. For all timepoints, the difference between the titres of ColorFlu-mCherry and ColorFlu-eGFP was not significant (Mann-Whitney U test, p>0.05). LLOQ = Lower limit of quantification.

**SUPPLEMENTARY DATA 2:**
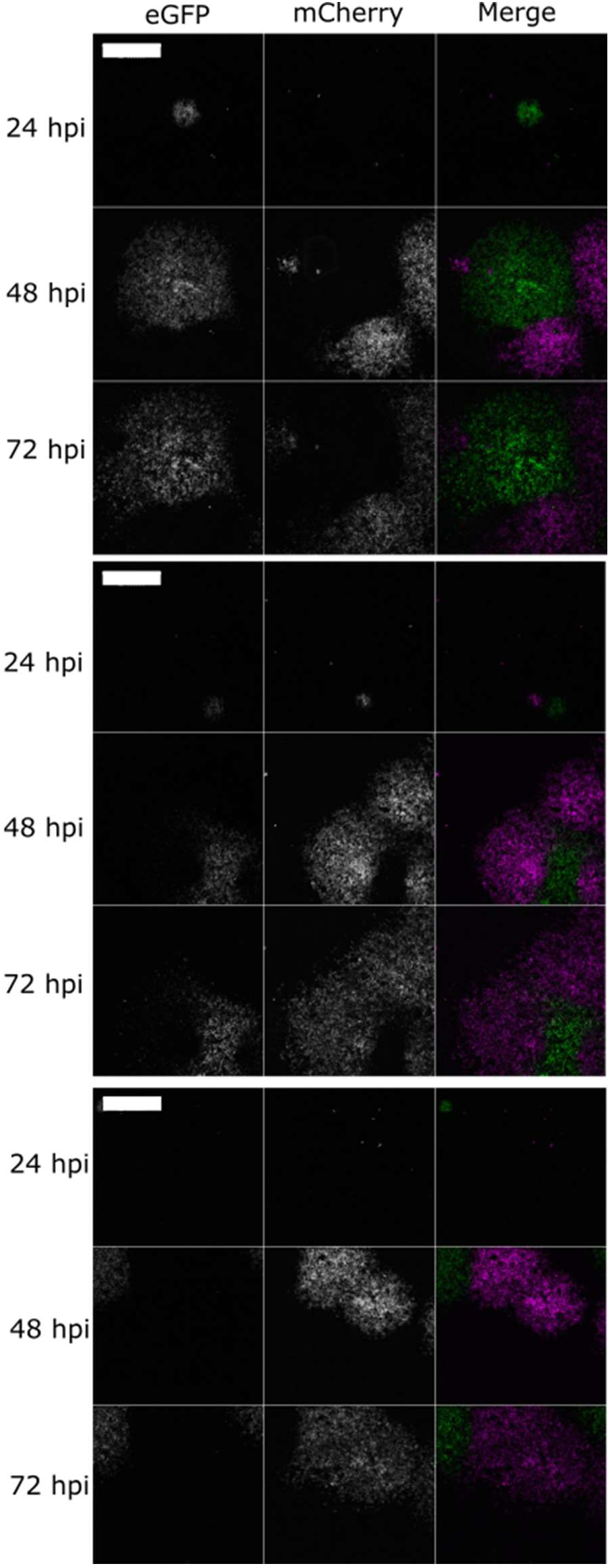
Further examples of superinfection exclusion limiting coinfection between distinct virus populations in vitro. Viruses were seeded onto monolayers of MDCK cells, overlayed with agarose and imaged every 24 hours. Images taken on Celigo fluorescent microscope. Scale bar = 2mm.

**SUPPLEMENTARY DATA 3:**
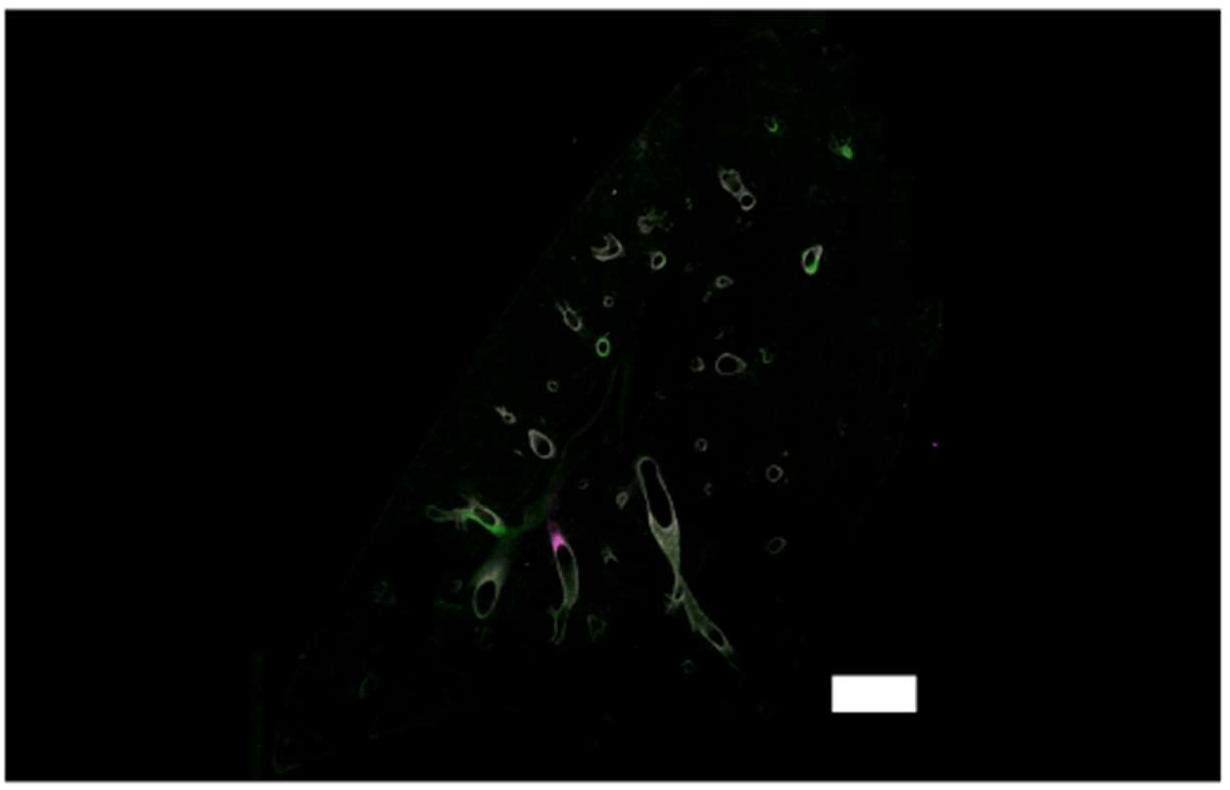
Initial mouse infection occurs in the bronchi. Whole lung images of ColorFlu infected mice 3 days post infection B57BL/6 mice were intranasally inoculated with mixtures of mCherry and eGFP expressing viruses (500 pfu of each virus). Lung sections at 3 days post infection were imaged using Zeiss LSM 800 using 20x objective lens. Scale bar = 1500µm.

**SUPPLEMENTARY DATA 4:**
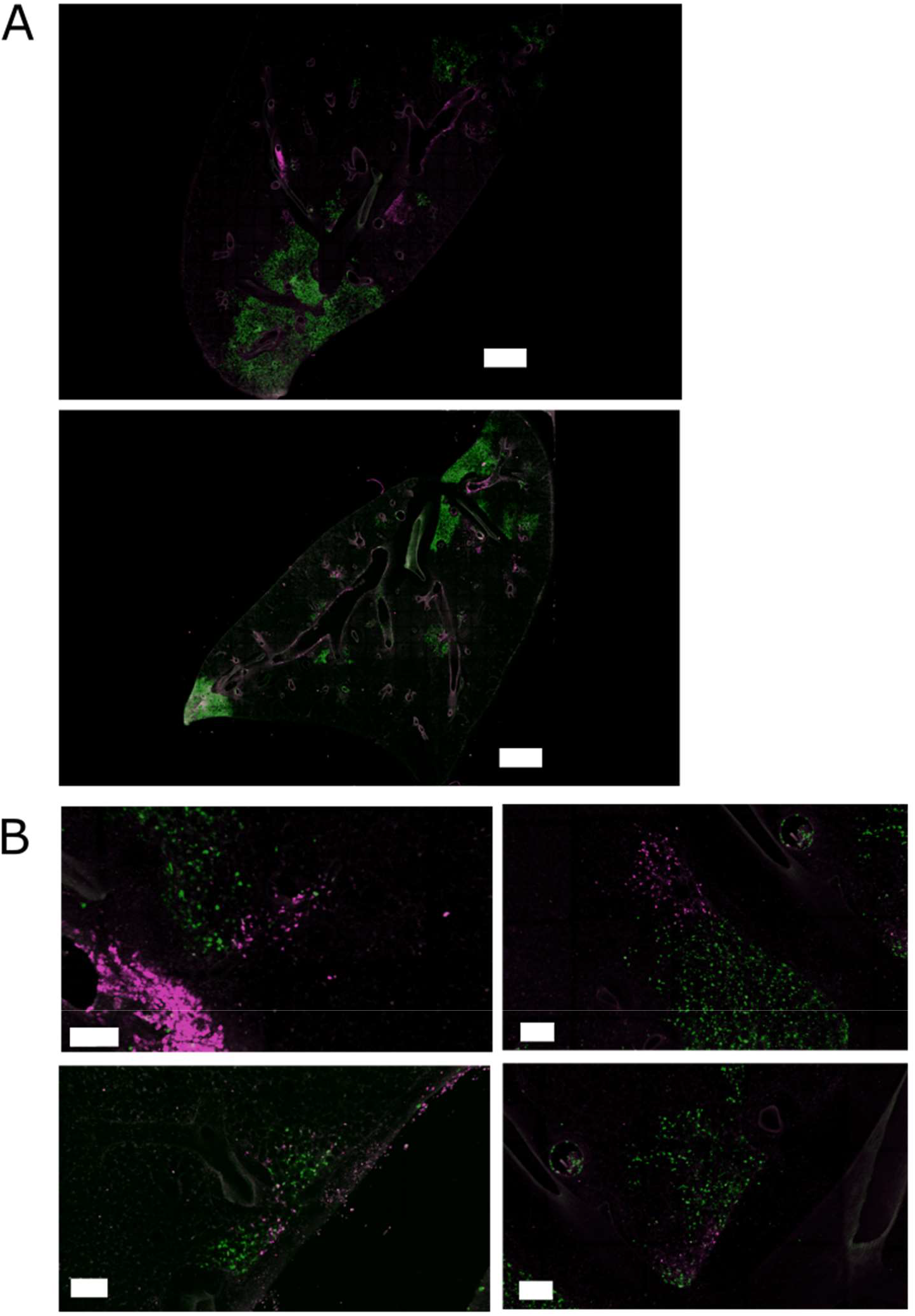
Further examples of superinfection exclusion limiting coinfection between distinct virus populations in vivo. **(A)** Confocal micrographs of whole lung slices from infected mice 6 dpi. B57BL/6 mice were intranasally inoculated with mixtures of mCherry and eGFP expressing viruses (500 pfu of each virus). Lung sections at 6 dpi were imaged using Zeiss LSM 800 using 20x objective lens. **(A)** Whole lung images. Scale bar = 1500µm **(B)** Enlarged images of infected lesions. Scale bar = 100µm.

**SUPPLEMENTARY MOVIE 1: Cells migrate inwards as infected plaques expand** Diluted mixtures of ColorFlu viruses were used to infect MDCK cells under agarose and observed over 72 hours in Zeiss Livecell observer microscope using a 20x objective lens.

## References

1. Paget J, Spreeuwenberg P, Charu V, Taylor RJ, Iuliano AD, Bresee J, et al. Global mortality associated with seasonal influenza epidemics: New burden estimates and predictors from the GLaMOR Project. J Glob Health. 2019;9(2):1–12.

2. Kim H, Webster RG, Webby RJ. Influenza Virus: Dealing with a Drifting and Shifting Pathogen. Viral Immunol. 2018 Mar;31(2):174–83.

3. Steel J, Lowen AC. Influenza A virus reassortment. Curr Top Microbiol Immunol. 2014;385:377– 401.

4. Brooke CB. Population Diversity and Collective Infection. 2017;91(22):1–13.

5. Vignuzzi M, López CB. Defective viral genomes are key drivers of the virus–host interaction. Nat Microbiol [Internet]. 2019;4(7):1075–87. Available from: http://dx.doi.org/10.1038/s41564-019-0465-y

6. Andreu-Moreno I, Sanjuán R. Collective Infection of Cells by Viral Aggregates Promotes Early Viral Proliferation and Reveals a Cellular-Level Allee Effect. Curr Biol. 2018 Oct 22;28(20):3212-3219.e4.

7. Davis AR, Nayak DP. Sequence relationships among defective interfering influenza viral RNAs. Proc Natl Acad Sci U S A. 1979;76(7):3092–6.

8. Ziegler CM, Botten JW. Defective Interfering Particles of Negative-Strand RNA Viruses. Trends Microbiol [Internet]. 2020;0–11. Available from: https://doi.org/10.1016/j.tim.2020.02.006

9. Brooke CB. Biological activities of “noninfectious” influenza A virus particles. Future Virol. 2014;9(1):41–51.

10. Brooke CB, Ince WL, Wrammert J, Ahmed R, Wilson PC, Bennink JR, et al. Most Influenza A Virions Fail To Express at Least One Essential Viral Protein. J Virol. 2013 Mar 15;87(6):3155–62.

11. Farrell A, Brooke C, Koelle K, Ke R. Coinfection of semi-infectious particles can contribute substantially to influenza infection dynamics. bioRxiv [Internet]. 2019;1–27. Available from: http://dx.doi.org/10.1101/547349

12. Hutchinson EC. Influenza Virus. Trends Microbiol. 2018;26(9):809–10.

13. Postnikova Y, Treshchalina A, Boravleva E, Gambaryan A, Ishmukhametov A, Matrosovich M, et al. Diversity and reassortment rate of influenza a viruses in wild ducks and gulls. Viruses. 2021;13(6):1–14.

14. Hill NJ, Hussein ITM, Davis KR, Ma EJ, Spivey TJ, Ramey AM, et al. Reassortment of Influenza A Viruses in Wild Birds in Alaska before H5 Clade 2.3.4.4 Outbreaks. Emerg Infect Dis [Internet]. 2017 Apr;23(4):654–7. Available from: https://pubmed.ncbi.nlm.nih.gov/28322698

15. Cui Y, Li Y, Li M, Zhao L, Wang D, Tian J, et al. Evolution and extensive reassortment of H5 influenza viruses isolated from wild birds in China over the past decade. Emerg Microbes Infect [Internet]. 2020 Jan 1;9(1):1793–803. Available from: https://doi.org/10.1080/22221751.2020.1797542

16. Jacobs NT, Onuoha NO, Antia A, Steel J, Antia R, Lowen AC. Incomplete influenza A virus genomes occur frequently but are readily complemented during localized viral spread. Nat Commun [Internet]. 2019;10(1). Available from: http://dx.doi.org/10.1038/s41467-019-11428-x

17. Marshall N, Priyamvada L, Ende Z, Steel J, Lowen AC. Influenza Virus Reassortment Occurs with High Frequency in the Absence of Segment Mismatch. PLoS Pathog. 2013 Jun;9(6).

18. Gallagher ME, Brooke CB, Ke R, Koelle K. Causes and consequences of spatial within-host viral spread. Viruses. 2018;10(11):1–23.

19. Manicassamy B, Manicassamy S, Belicha-villanueva A, Pisanelli G, Pulendran B. Analysis of in vivo dynamics of in fl uenza virus infection in mice using a GFP reporter virus. 2010;107(25):11531–6.

20. Brand JMA Van Den, Stittelaar KJ, Amerongen G Van, Reperant L, De L, Osterhaus Adme, et al. Comparison of Temporal and Spatial Dynamics of Seasonal H3N2, Pandemic H1N1 and Highly Pathogenic Avian Influenza H5N1 Virus Infections in Ferrets. 2012;7(8).

21. Guarner J, Shieh WJ, Dawson J, Subbarao K, Shaw M, Ferebee T, et al. Immunohistochemical and in situ hybridization studies of influenza A virus infection in human lungs. Am J Clin Pathol. 2000 Aug;114(2):227–33.

22. Phipps KL, Ganti K, Jacobs NT, Lee C, Carnaccini S, White MC, et al. Collective interactions augment influenza A virus replication in a host-dependent manner. Nat Microbiol [Internet]. Available from: http://dx.doi.org/10.1038/s41564-020-0749-2

23. Zou G, Zhang B, Lim P-Y, Yuan Z, Bernard KA, Shi P-Y. Exclusion of West Nile Virus Superinfection through RNA Replication. J Virol. 2009;83(22):11765–76.

24. Schaller T, Appel N, Koutsoudakis G, Kallis S, Lohmann V, Pietschmann T, et al. Analysis of Hepatitis C Virus Superinfection Exclusion by Using Novel Fluorochrome Gene-Tagged Viral Genomes. J Virol. 2007;81(9):4591–603.

25. Laliberte JP, Moss B. A Novel Mode of Poxvirus Superinfection Exclusion That Prevents Fusion of the Lipid Bilayers of Viral and Cellular Membranes. J Virol. 2014;88(17):9751–68.

26. Zhang XF, Sun R, Guo Q, Zhang S, Meulia T, Halfmann R, et al. A self-perpetuating repressive state of a viral replication protein blocks superinfection by the same virus. PLoS Pathog. 2017;13(3):1–24.

27. McAllister WT, Barrett CL. Superinfection exclusion by bacteriophage T7. J Virol. 1977;24(2):709–11.

28. Kumar N, Sharma S, Barua S, Tripathi BN, Rouse BT. Virological and Immunological Outcomes of Coinfections. Clin Microbiol Rev. 2018 Oct;31(4).

29. Dou D, Hernández-Neuta I, Wang H, Östbye H, Qian X, Thiele S, et al. Analysis of IAV Replication and Co-infection Dynamics by a Versatile RNA Viral Genome Labeling Method. Cell Rep. 2017;20(1):251–63.

30. Greenbaum BD, Li OTW, Poon LLM, Levine AJ, Rabadan R. Viral reassortment as an information exchange between viral segments. Proc Natl Acad Sci U S A. 2012;109(9):3341–6.

31. White MC, Lowen AC. Implications of segment mismatch for influenza A virus evolution. J Gen Virol. 2018;99(1):3–16.

32. Fukuyama S, Katsura H, Zhao D, Ozawa M, Ando T, Shoemaker JE, et al. Multi-spectral fluorescent reporter influenza viruses (Color-flu) as powerful tools for in vivo studies. Nat Commun. 2015;6.

33. Bodewes R, Nieuwkoop NJ, Verburgh RJ, Fouchier RAM, Osterhaus Adme, Rimmelzwaan GF. Use of influenza A viruses expressing reporter genes to assess the frequency of double infections in vitro. J Gen Virol. 2012;93(8):1645–8.

34. Dou D, Revol R, Östbye H, Wang H, Daniels R. Influenza A virus cell entry, replication, virion assembly and movement. Vol. 9, Frontiers in Immunology. Frontiers Media S.A.; 2018.

35. Sun J, Brooke CB. Influenza A Virus Superinfection Potential Is Regulated by Viral Genomic Heterogeneity. MBio. 2018;9(5):1–13.

36. Vester D, Lagoda A, Hoffmann D, Seitz C, Heldt S, Bettenbrock K, et al. Real-time RT-qPCR assay for the analysis of human influenza A virus transcription and replication dynamics. J Virol Methods. 2010;168(1–2):63–71.

37. Hatada E, Hasegawa M, Mukaigawa J, Shimizu K, Fukuda R. Control of influenza virus gene expression: Quantitative analysis of each viral RNA species in infected cells. J Biochem. 1989;105(4):537–46.

38. Hutchinson EC, Yamauchi Y. Understanding Influenza. In: Yamauchi Y, editor. Influenza Virus: Methods and Protocols [Internet]. New York, NY: Springer New York; 2018. p. 1–21. Available from: https://doi.org/10.1007/978-1-4939-8678-1_1

39. Frensing T, Kupke SY, Bachmann M, Fritzsche S, Gallo-ramirez LE, Reichl U. Influenza virus intracellular replication dynamics, release kinetics, and particle morphology during propagation in MDCK cells. Appl Microbiol Biotechnol [Internet]. 2016;7181–92. Available from: http://dx.doi.org/10.1007/s00253-016-7542-4

40. Fonville JM, Marshall N, Tao H, Steel J, Lowen AC. Influenza Virus Reassortment Is Enhanced by Semi-infectious Particles but Can Be Suppressed by Defective Interfering Particles. PLoS Pathog. 2015;11(10).

41. Webster RG, Bean WJ, Gorman OT, Chambers TM, Kawaoka Y. Evolution and ecology of influenza A viruses. Microbiol Rev. 1992 Mar;56(1):152–79.

42. Chen LM, Davis CT, Zhou H, Cox NJ, Donis RO. Genetic compatibility and virulence of reassortants derived from contemporary avian H5N1 and human H3N2 influenza A viruses. PLoS Pathog. 2008;4(5):1–11.

43. Kilbourne ED. Future influenza vaccines and the use of genetic recombinants. Bull World Health Organ. 1969;41(3):643–5.

44. Nakatsu S, Sagara H, Sakai-Tagawa Y, Sugaya N, Noda T, Kawaoka Y. Complete and incomplete genome packaging of influenza A and B viruses. MBio. 2016;7(5):1–7.

45. Diefenbacher M, Sun J, Brooke CB. The parts are greater than the whole: the role of semi-infectious particles in influenza A virus biology. Curr Opin Virol [Internet]. 2018;33:42–6. Available from: https://doi.org/10.1016/j.coviro.2018.07.002

46. Hunter M, Fusco D. Superinfection exclusion: A viral strategy with short-term benefits and long-term drawbacks. PLOS Comput Biol [Internet]. 2022 May 10;18(5):e1010125. Available from: https://doi.org/10.1371/journal.pcbi.1010125

47. Rezelj V V., Levi LI, Vignuzzi M. The defective component of viral populations. Curr Opin Virol [Internet]. 2018;33:74–80. Available from: https://doi.org/10.1016/j.coviro.2018.07.014

48. Huang I-C, Li W, Sui J, Marasco W, Choe H, Farzan M. Influenza A Virus Neuraminidase Limits Viral Superinfection. J Virol. 2008;82(10):4834–43.

49. Gaush CR, Smith TF. Replication and plaque assay of influenza virus in an established line of canine kidney cells. Appl Microbiol. 1968;16(4):588–94.

50. Schindelin J, Arganda-Carreras I, Frise E, Kaynig V, Longair M, Pietzsch T, et al. Fiji: An open-source platform for biological-image analysis. Nat Methods. 2012;9(7):676–82.

51. Figliozzi RW, Chen F, Chi A, Hsia SCV. Using the inverse Poisson distribution to calculate multiplicity of infection and viral replication by a high-throughput fluorescent imaging system. Virol Sin. 2016;31(2):180–3.

